# Analysis of the gut and gill microbiome of resistant and susceptible lines of rainbow trout (*Oncorhynchus mykiss*)

**DOI:** 10.1101/420018

**Authors:** Ryan M. Brown, Gregory D. Wiens, Irene Salinas

## Abstract

Commensal microorganisms present at mucosal surfaces play a vital role in protecting the host organism from bacterial infection. There are multiple factors that contribute to selecting for the microbiome, key of which are host genetics. *Flavobacterium psychrophillum*, the causative agent of Bacterial Cold Water Disease in salmonids, accounts for acute losses in wild and farmed Rainbow Trout (*Oncorhynchus mykiss*). The U.S. National Center for Cool and Cold Water Aquaculture has used family-based selective breeding to generate a line of rainbow trout with enhanced resistance to *F. psychrophilum*. The goal of this study is to determine whether selective breeding impacts the gut and gill microbiome of the *F. psychrophilum*-resistant as compared to a background matched susceptible trout line. Mid-gut and gill samples were collected from juvenile fish (mean bwt 118g) and microbial diversity assessed by 16S rDNA amplicon sequencing. Results indicate that alpha diversity was significantly higher in the mid-gut of the susceptible line compared to the resistant line, while no significant differences in alpha diversity were observed in the gills. *Mycoplasma sp*. was the dominant taxon in the mid-gut of both groups, although it was present at a decreased abundance in the susceptible line. We also observed an increased abundance of taxa that could potentially be pathogenic in the susceptible line, including *Brevinema sp*. and Enterobacteriaceae members. Within the gills, both lines exhibited similar microbial profiles, with *Candidatus Branchiomonas* being the dominant taxon. Together, these results suggest that selectively bred *Flavobacterium psychrophillum*-resistant trout may harness a more resilient gut microbiome, attributing to the disease resistant phenotype, providing a framework for future experiments.

## 1. Introduction

The microbiome has well established roles in pathogen exclusion and host immunity, including systemic and mucosal innate and adaptive immune responses and development of the immune system [1–3]. There is strong evidence to support a role of host genetics in the selection of the gut microbiome in humans and other mammals [4–6], although this has not been well characterized in fish. The host microbial composition is also shaped by other factors including environment, diet, and disease [7–10]. To begin to disentangle the contribution of host genetics and environmental factors shaping the fish microbiome, here we utilize a rainbow trout model in which two genetic lines of rainbow trout have been established by selective breeding that differ in susceptibility to a common environmental gram negative pathogen, *Flavobacterium psychrophilum*.

*Flavobacterium psychrophilum* is the causative agent of Bacterial Cold Water Disease (BCWD), which is a major concern in the United States aquaculture industry affecting a range of cold-water fish species, including the commercially relevant rainbow trout (*Oncorhynchus mykiss*). *F. psychrophillum* is a mucosal pathogen that typically infects the skin and gills of fish [11]. Symptoms of BCWD in developed fish include necrosis of the caudal region, skin lesions, eroded fin tips, and loss of appetite. *F. psychrophilum* has a more pronounced effect on young fry, a condition referred to as rainbow trout fry syndrome. Rainbow trout fry syndrome is responsible for acute losses in trout farms worldwide, as the associated mortality rate is reported to be greater than 50% [12]. BCWD is becoming an increasingly difficult disease to treat, as *F. psychrophilum* strains have developed resistance to several commonly used antibiotics [13–15], and there is currently no commercially available licensed vaccine.

The National Center for Cool and Cold Water Aquaculture (NCCCWA) utilized family-based selective breeding to develop two distinctive genetic lines of rainbow trout that confer enhanced resistance (ARS-Fp-R), or susceptibility (ARS-Fp-S) to the pathogen *F. psychrophilum* [16]. Enhanced resistance to *F. psychrophilum*-induced mortality in the ARS-Fp-R line has been described, both in the laboratory setting and on trout farms [17,18]. Previous studies have begun to investigate possible host mechanisms that attribute to enhanced resistance. For instance, a strong correlation between resistance to *F. psychrophilum* and increased spleen size has been described, although this relation does not appear to translate to other common fish pathogens, such as *Yersinia ruckeri* [19]. Additionally, whole-body transcriptome analysis has identified numerous acute phase proteins and inflammatory cytokines that are differentially expressed in each line following challenge with *F. psychrophilum* [20]. Further work is needed to better characterize the mechanism(s) by which enhanced resistance is achieved in the ARS-Fp-R line.

In this paper, we begin to investigate the hypothesis that the higher innate disease resistance of the ARS-Fp-R line is due to differences in the microbial composition at host mucosal surfaces. Using 16S rDNA amplicon sequencing, we determine and compare the mid-gut and gill microbiomes of rainbow trout lines selectively bred for *F. psychrophilum* resistance and susceptibility.

## 2. Materials and Methods

### 2.1. Animals and sampling

The Institutional Animal Care and Use Committee (Leetown, WV) reviewed and approved all animal husbandry practices and disease challenge protocols per standards set forth in the USDA, ARS Policies and Procedures 130.4.v.3 titled ‘Institutional Animal Care and Use 84 Committee’. Fish used in these experiments were from the 2017 Year Class and maintained as specific pathogen free as determined by biannual testing as previously described [17]. A total of 33 single sire-dam matings contributed to the pool of ARS-Fp-R line fish and 31 matings contributed to the pool of ARS-Fp-S line. The disease resistant phenotype of each genetic line was evaluated at two time-points, 75 and 276 days post-hatch. Fish were challenged with *F. psychrophilum* strain CSF259-93 and survival recorded over 21 days as previously described [19]. Mean fish body weight at the first evaluation was 1.9 g and a total of 120 ARS-Fp-R and 119 ARS-Fp-S fish (n=3 tanks per line) were challenged by intraperitoneal injection with a dose of 1.4E+07 CFU g^−1^ in a total volume of 50 μL using a 26g needle fitted onto an Eppendorf repeating pipette. Mean body weight at the second evaluation was 194 g and a total of 70 ARS-Fp-R and 70 ARS-Fp-S fish (n=2 tanks per line) were challenged by intramuscular injection with a dose of 3.5E+06 CFU g^−1^ body weight in a total volume of 50 μL using a 26g needle fitted onto an Eppendorf repeating pipette. The fish utilized in the second experiment were part of a larger study evaluating experimental vaccination and these fish had been sham vaccinated with PBS 35 days prior to challenge. In both challenges, *F. psychrophilum* was isolated from mortalities and confirmed by PCR genotyping.

At the time of microbiome sampling, fish were reared under two different tank conditions as described in Supplemental data file 1. Ten fish from each tank system per genetic line were euthanized using 200 mg L^-1^ MS222 for 5 minutes. Each fish was photographed, weighted, gill tissue sampled, intestinal mid-gut sampled, and spleen weighed within 30 minutes of euthanasia. Sample were placed in SLB [21] on ice and then moved to storage at−80° C. Instruments were cleaned between each fish and gloves changed between tank groups. Three control tubes containing SLB alone were included as negative controls.

### 2.2. DNA Extraction, 16S rDNA PCR Amplification, and Sequencing

Whole genomic DNA was extracted from skin and gill samples by first lysing tissue samples using sterile 3 mm tungsten beads (Qiagen) and Qiagen TissueLyser II. Next, using the cetyltrimethylammonium bromide method as previously described [21], DNA was isolated and suspended in 50 μL RNase and DNase free molecular biology grade water. DNA concentration and purity was then assessed using a Nanodrop ND 1000 (Thermo Scientific).

Bacterial DNA was then replicated by PCR using Illumina adapter fused primers targeting the V1-V3 region of the prokaryotic 16S rDNA gene. The primer sequences were as follows: 28F 5’-GAGTTTGATCNTGGCTCAG-3’ and 519R 5’GTNTTACNGCGGCKGCTG-3’ (where N = any nucleotide, and K = T or G). DNA samples were diluted 1:10 or 1:100, and Quantabio 5PRIME HotMasterMix was used. 16S amplicons were generated using the following conditions: 94° C for 90s; 33 cycles of 94° C for 30s, 52° C for 30s, 72° C for 90s; and a final extension of 72° C for 7 min A positive control of a verified 16S V1-V3 amplicon, and a negative control of molecular biology grade water was included in every PCR reaction. In addition, we included a mock community positive control with each sequencing run, which consisted of equal DNA amounts of 7 different bacterial isolates previously cloned and a negative control that consisted of SLB handled in the same way as the rest of the tubes during sampling to which no tissue was added. Amplicons were purified using the Axygen AxyPrep Mag PCR Clean-up Kit (Thermo Scientific), and eluted into 30 μL molecular biology grade water. Unique oligonucleotide barcodes were ligated to the 5′ and 3′ ends of each sample, as well as the Nextera adaptor sequences, using the Nextera XT Index Kit v2 set A (Illumina). DNA concentrations were quantified using a Qubit, and normalized to a concentration of 200 ng/uL for DNA library pooling. Pooled samples were cleaned once more using the Axygen PCR clean-up kit before being sent off for sequencing. Sequencing was performed on the Illumina Miseq platform using the MiSeq Reagent Kit v3 (600 cycle) at the Clinical and Translational Sciences Center at the University of New Mexico Health Sciences Center.

### 2.3. Data Analysis and statistics

Differences in survival between genetic lines were determined using the product limit method of Kaplan and Meier and calculations were performed using GraphPad v4.0 software. Log-rank (Mantel-Cox) test was used to compare survival curves. Initial sequence data was analyzed using the latest version of Quantitative Insights Into Microbial Ecology 2 (Qiime2 v2018.6) [22]. Demultiplexed sequence reads were quality filtered using DADA2 [23]. Samples were then rarefied to a sampling depth of 12,603 sequences per sample for mid-gut reads, and 2020 sequences per sample for gill reads. Amplicon sequence variants (ASVs) were picked by aligning to the latest Silva 16S database (version 132). Core diversity metrics were analyzed, including number of ASVs and Shannon’s diversity index for alpha diversity, and PERMANOVA for beta diversity. Nonmetric multidimensional scaling and generation of heat maps were performed in RStudio [24] using the phyloseq package [25]. Random forest modeling was performed in Qiime2. Differential abundance testing was performed in Qiiime2 using ANCOM [26], as well as in RStudio using the DESeq2 package [27]. For all statistical analyses, fish were split into groups based on ‘Treatment’ (ARS-Fp-R, ARS-Fp-S) or ‘Tank Treatment’.

## 3. Results

### 3.1. Phenotype Confirmation of Disease Resistance/Susceptibility

The relative phenotype of the two genetic lines was evaluated at time-points either before or after microbiome sampling. At both time points, the survival of the ARS-Fp-R genetic line was significantly higher (*P*<0.001) than the ARS-Fp-S line and consistent with estimated mid-point breeding values. In the first evaluation, a total of 3/120 (3%) ARS-Fp-R line fish died compared to 82/119 (69%) ARS-Fp-S line fish. In the second evaluation, 2/70 (3%) ARS-Fp-R fish died, while 58/70 (83%) ARS-Fp-S line fish died within the 21-day challenge period.

### 3.2. High throughput sequencing analysis

A total number of 3,598,038 raw reads were obtained from all mid-gut samples. After merging paired ends, quality filtering, and removal of chimeric reads with DADA2, as well as filtering out non-specific trout genomic reads, a total of 1,413,104 reads remained, with a mean of 35,328 reads per sample. Samples were rarified to a sample depth of 12,031, which excluded two ARS-Fp-R samples and two ARS-Fp-S samples. The sample size after rarefaction was n=18 for both the ARS-Fp-R and ARS-Fp-S lines.

Gill sample sequencing produced a total of 4,646,971 raw reads. Quality filtering in DADA2 and removal of non-specific reads retained 333,281 reads, with a mean of 13,331 reads per sample. Samples were rarified to 2010 reads. The sample size after rarefaction was n=11 for ARS-Fp-R and n=14 for ARS-Fp-S.

### 3.3. Resistant and susceptible lines display significant differences in alpha diversity in the mid-gut, but not the gills

Comparison of alpha diversity metrics obtained from the mid-gut showed significantly lower measures of gut microbial community richness (Observed ASVs), as well as Shannon’s diversity index in the ARS-Fp-R line compared to the ARS-Fp-S line (Fig 1A and 1B). There were a total of 15 ASVs reported in the mid-gut of the resistant line, and 29 in the mid-gut of the susceptible line. In the gills, there were no significant differences in alpha diversity between lines with a total of 57 ASVs found in the gills of the resistant line, and 50 in the susceptible line (Fig 1C and 1D).

**Fig 1.**
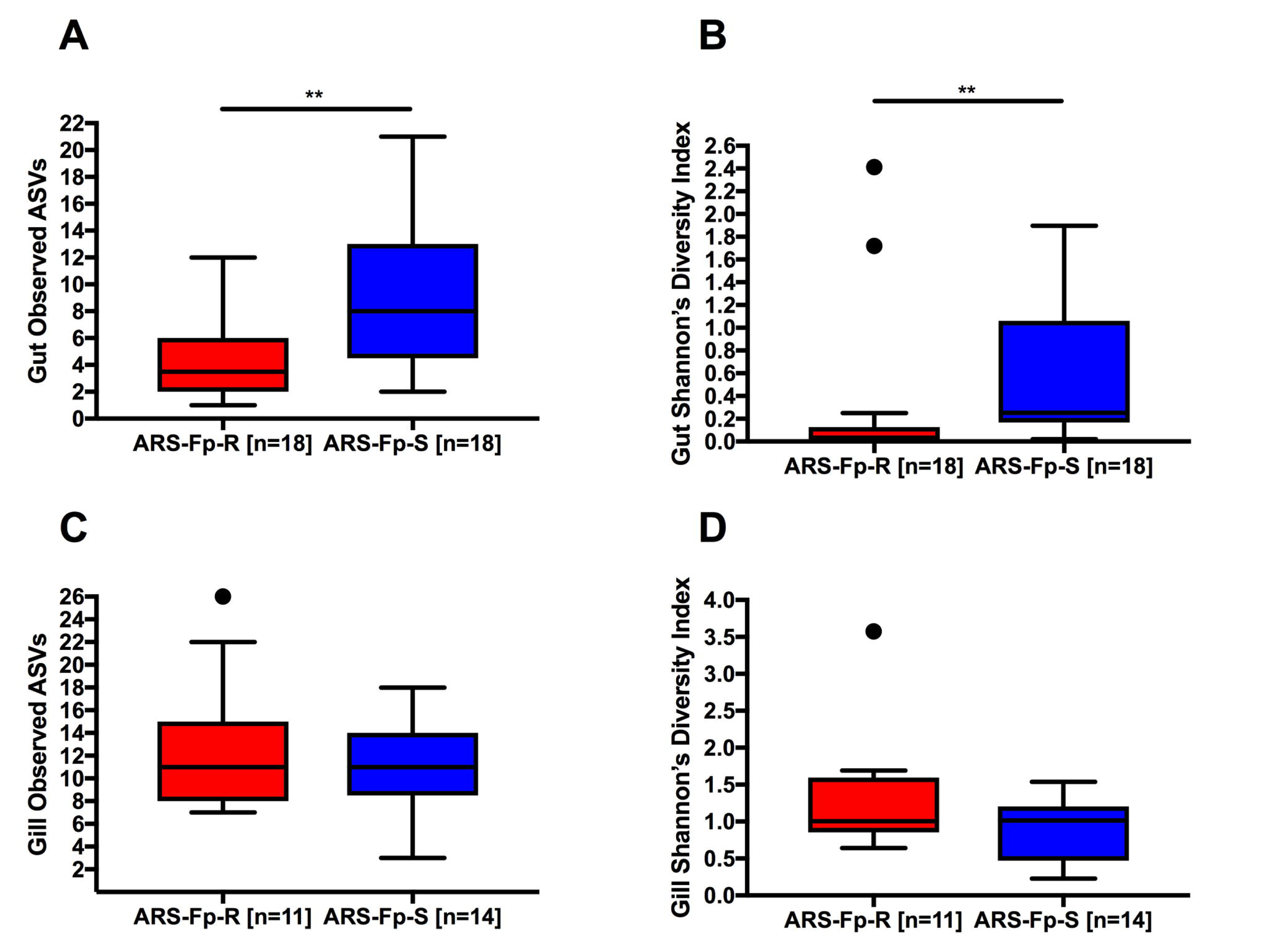
Comparison of alpha diversity metrics for the mid-gut and gill microbiome of ARS-Fp-R and ARS-Fp-S trout. (A) Total number of observed ASVs in the mid-gut. (B) Shannon’s diversity index in the mid-gut. (C) Total number of observed ASVs in the gills. (D) Shannon’s diversity index in the gills. ** indicates statistically significant differences p<0.01.

### 3.4. Beta diversity analysis suggests possible tank effect on the gut microbiome

We assessed the microbial diversity between different treatments, as well as between tanks by performing Nonmetric Multidimensional Scaling (NMDS) using the Bray Curtis distance metric. This ordination showed a discrete grouping of the two tanks containing the ARS-Fp-S line, while the two tanks containing the ARS-Fp-R line were more tightly clustered (Fig 2A). PERMANOVA analysis [28] identified “Treatment” (P value = 0.02) and “Tank” (P value = 0.048 for ARS-Fp-R, P value = 0.001 for ARS-Fp-S) as significant determinants of the mid-gut microbial community composition. In the gills, NMDS ordination showed a similar pattern to that found in the gut, where fish from tank 26 of the susceptible line clustered tightly together while fish from tank 12 showed greater variability. Meanwhile, there was no clear separation between individuals held in separate tanks of the resistant line (Fig 2B). PERMANOVA identified only tank housing within the ARS-Fp-S line (P value = 0.004) as a significant factor in determining gill microbial communities.

**Fig 2.**
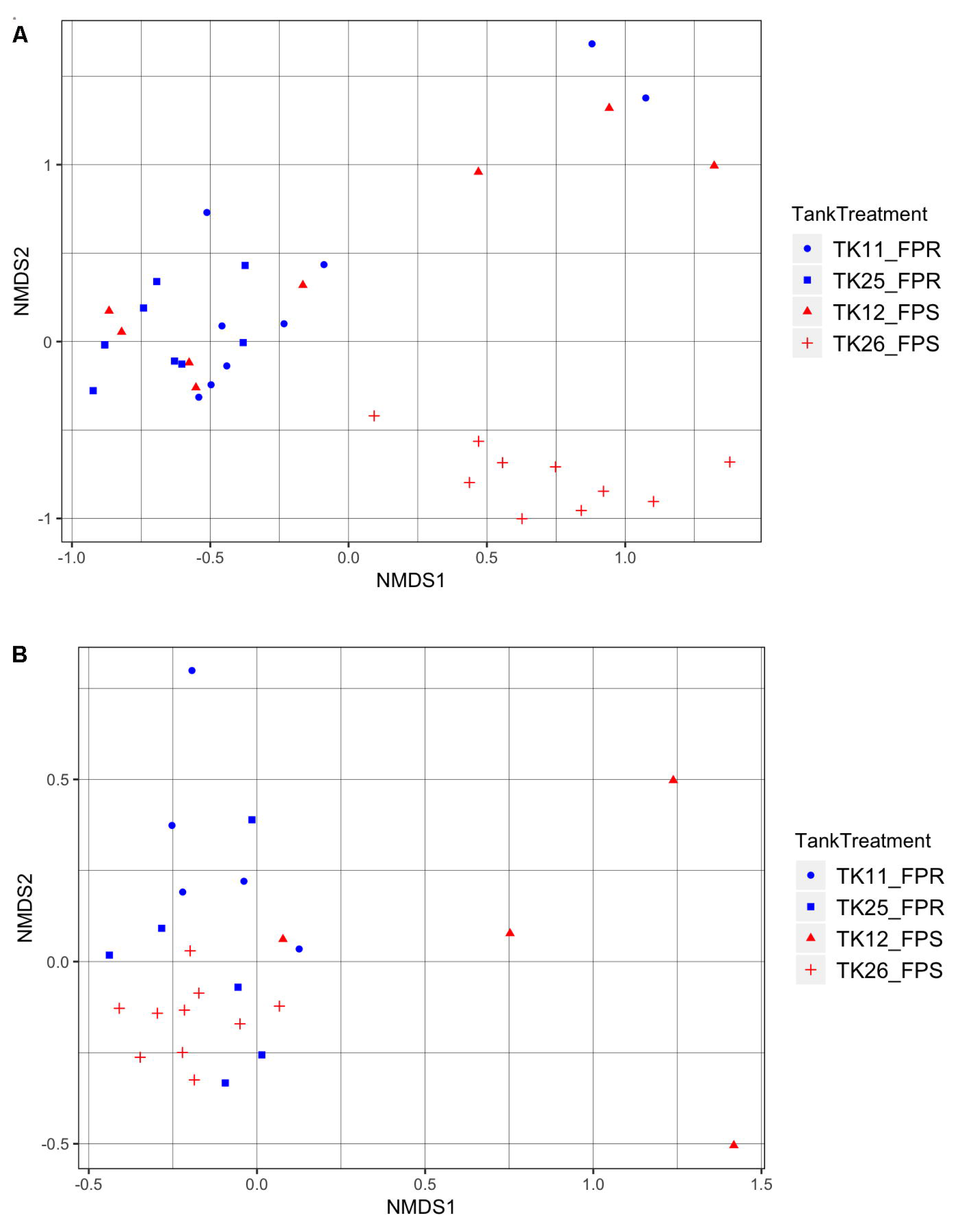
NMDS ordination plots. NMDS ordination performed using Bray Curtis distance matrix of the (A) mid-gut and (B) gill microbiome of ARS-Fp-R and ARS-Fp-S trout.

### 3.5. Gut microbial community composition

A total of eight different phyla were identified in the mid-gut across both lines, although only four of these were represented over 1%, including Tenericutes, Spirochaetes, Proteobacteria, and Firmicutes. Tenericutes composed the vast majority of the mid-gut microbiome of both lines, constituting 81% of the total microbial diversity in the susceptible line and 89% in the resistant line (Fig 3A). At the genus level, all Tenericutes reads were identified as *Mycoplasma sp*. The genus *Brevinema sp.* showed greater abundance in the ARS-Fp-S (8.5%) compared to the ARS-Fp-R line (4.8%). Similarly, members of the family Enterobacteriaceae had greater abundance in the ARS-Fp-S line than the ARS-Fp-R line (2.2% and 0%, respectively); however, these differences were not significant. Differential abundance testing with ANCOM revealed three genera that were differentially abundant in the mid-gut of both trout lines; including *Hydrotalea sp., Paenibacillus sp., and Variovorax sp.* This was replicated by differential abundance testing using DESeq2, an R package originally developed for differential expression analysis in RNA-seq data, but has been proposed to be used in microbiome studies as well [29,30].

**Fig 3.**
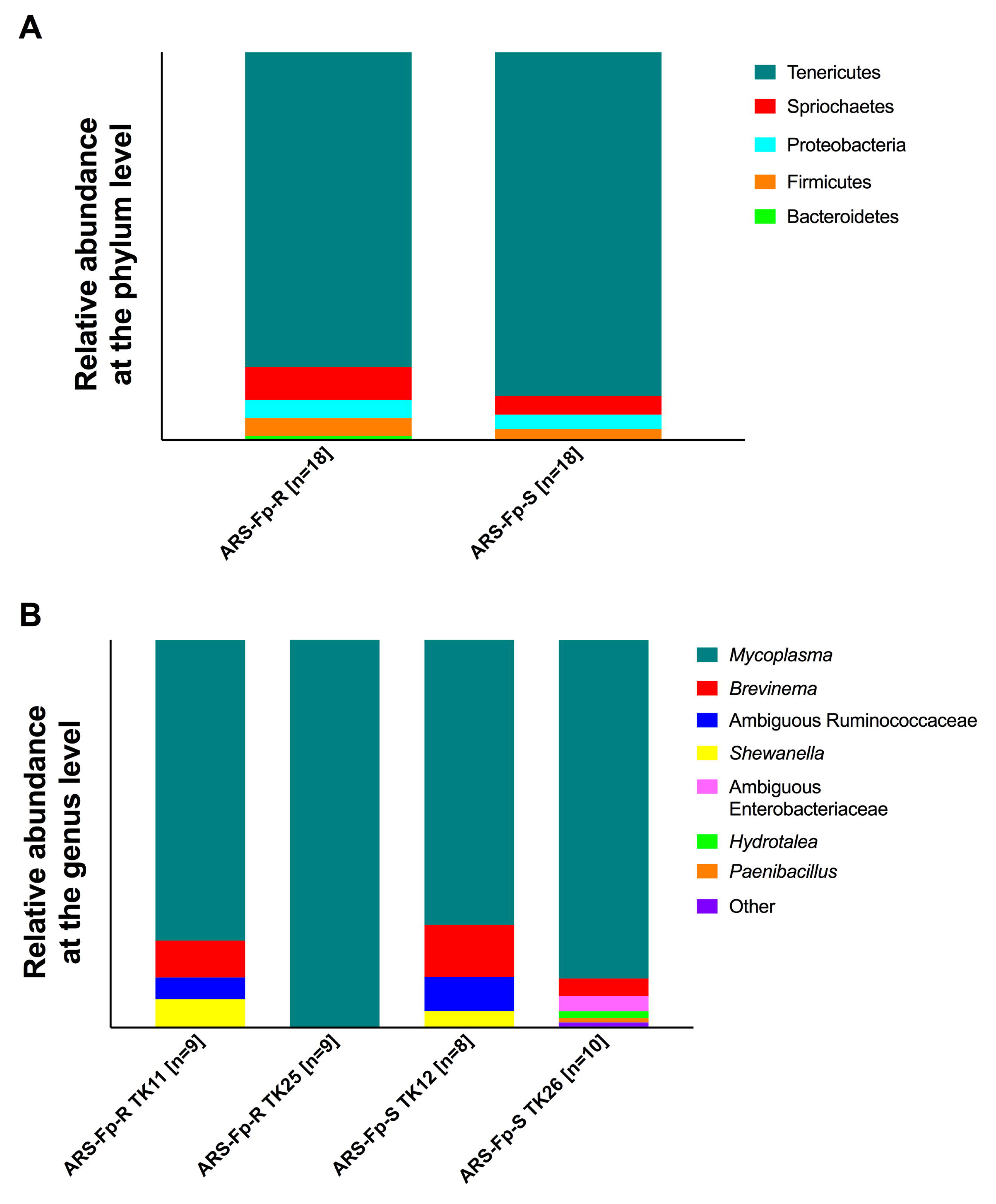
Relative microbial composition in the gut of each line. (A) Relative abundance of ASVs at the phylum level for each line. (B) Relative abundance of ASVs for each tank the two lines were divided into.

The relative distribution of ASVs at the genus level was notably different between tanks of the same line (Fig 3B). Potential opportunistic taxa such as *Brevinema sp.* and Ambiguous Enterobacteriaceae were present in tanks 11 and 12, but were not identified in either tank 25 or 26. This trend is further shown in a heatmap representing the top 25 ASVs observed in each sample (Fig 4). Two of the tanks that were in close proximity to one another displayed similar microbial profiles, despite holding different lines. This was not the case for tanks 25 and 26, as tank 26 displayed a microbial profile different from all other tanks since it contained ASVs representative of *Enterobacterieaceae, Hydrotalea sp., Paenibacillus sp*, and *Variovorax sp*.

**Fig 4.**
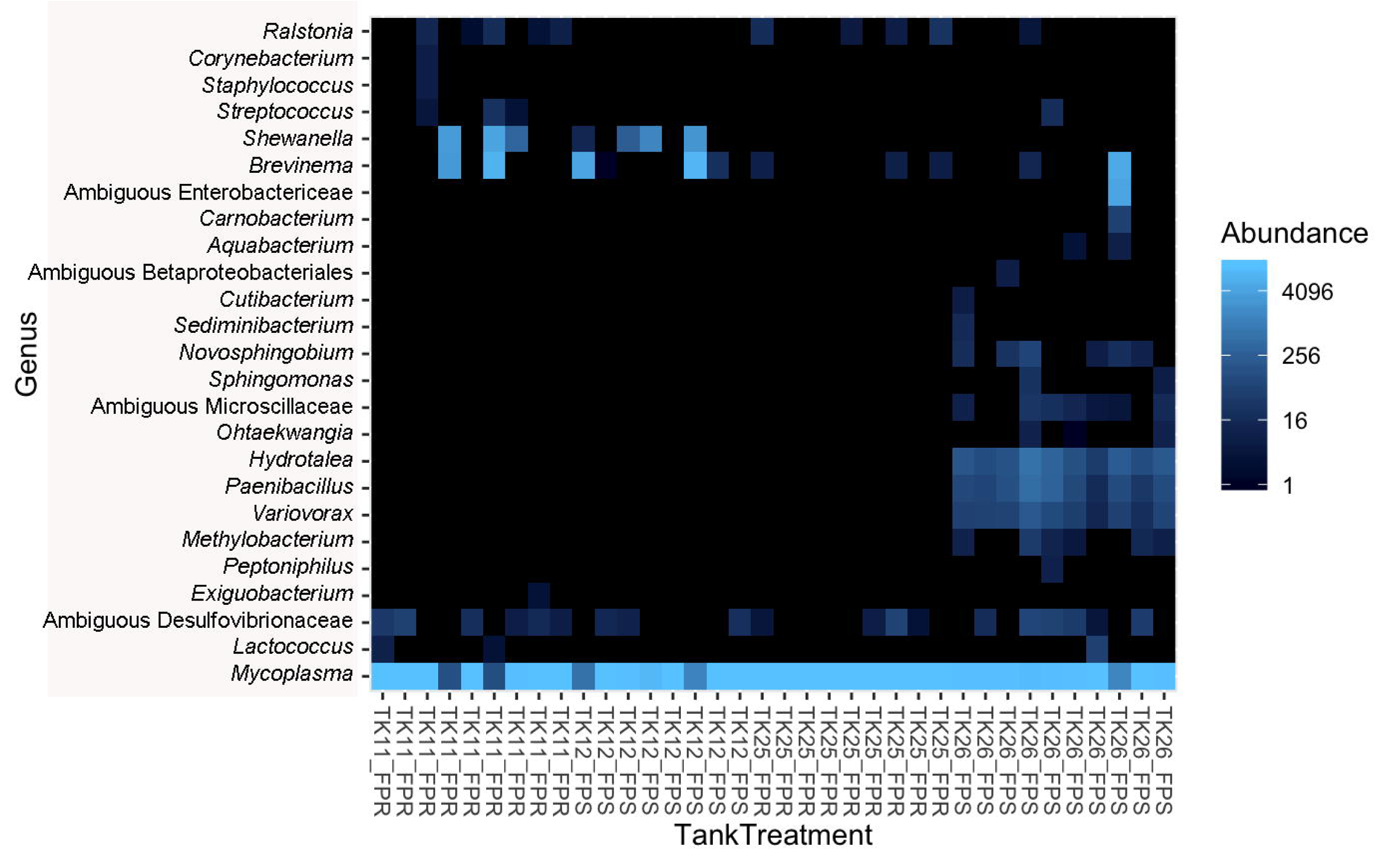
Heatmap representing the top 25 ASVs present in the mid-gut of ARS-Fp-R and ARS-Fp-S trout. Each column represents one individual. Each row represents one ASV.

### 3.6. Gill microbial community composition

A total of nine different phyla were present in the trout gills across both lines. Five of these were represented at abundance greater than 1%, including Proteobacteria, Tenericutes, Spirochaetes, Bacteriodetes, and Firmicutes. Proteobacteria was the most abundant phylum in both groups, representing 85% of all bacterial diversity in the ARS-Fp-S line and 95% in the ARS-Fp-R line (Fig 5A). At the genus level, most Proteobacteria reads were identified as *Candidatus Branchiomonas*. This taxon constituted 74% of all diversity in the gills of the susceptible line and 85% in the resistant line. We identified trace amounts of *Flavobacterium sp*. in both lines, as this taxon constituted 0.16 % of the ARS-Fp-S line and 1.1% of the ARS-Fp-R line. No ASVs were differentially abundant in the gill microbial community of both groups through ANCOM or DEseq2 analyses.

**Fig 5.**
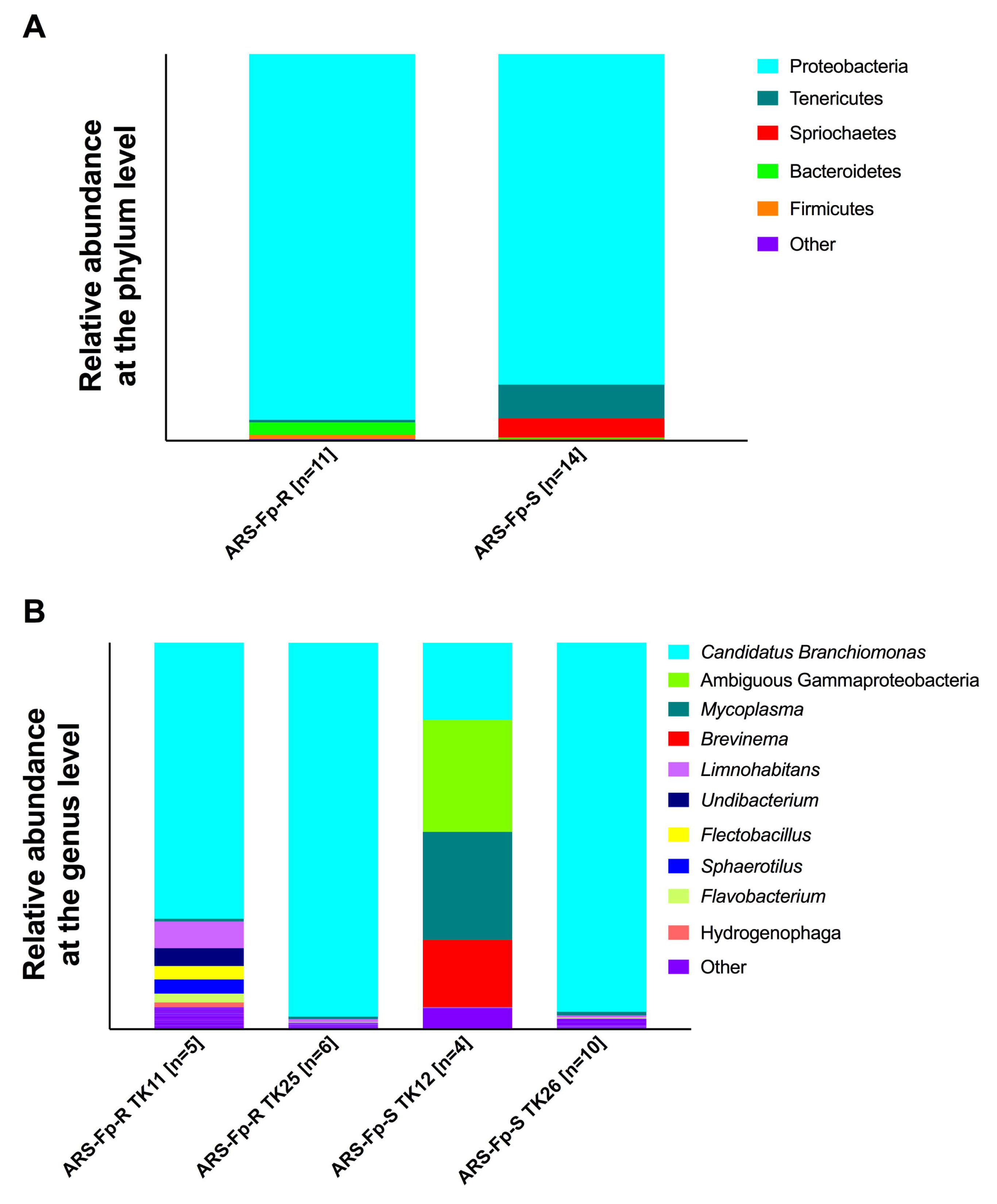
Microbial community composition of the gills of ARS-Fp-R and ARS-Fp-S trout. (A) Relative abundance of ASVs at the phylum level for each line. (B) Relative abundance of ASVs for tank the two lines were divided into.

We observed discernable differences in the microbial community composition of fish of the same line housed in different tanks (Fig 5B). For example, *Candidatus Branchiomonas* was present at levels above 95% in tanks 25 (ARS-Fp-R) and 26 (ARS-Fp-S), whereas it constituted 71% of tank 11 (ARS-Fp-R) and 20% of tank 12 (ARS-Fp-S). The aforementioned potential opportunistic pathogens *Brevinema sp*. was only identified in tank 12. A heatmap of the top 30 ASVs in each sample (Fig 6) shows signatures of *Brevinema sp*. in fish from tank 12, as well as a reduced abundance of *Candidatus Branchiomonas* in this tank compared to all others.

**Fig 6.**
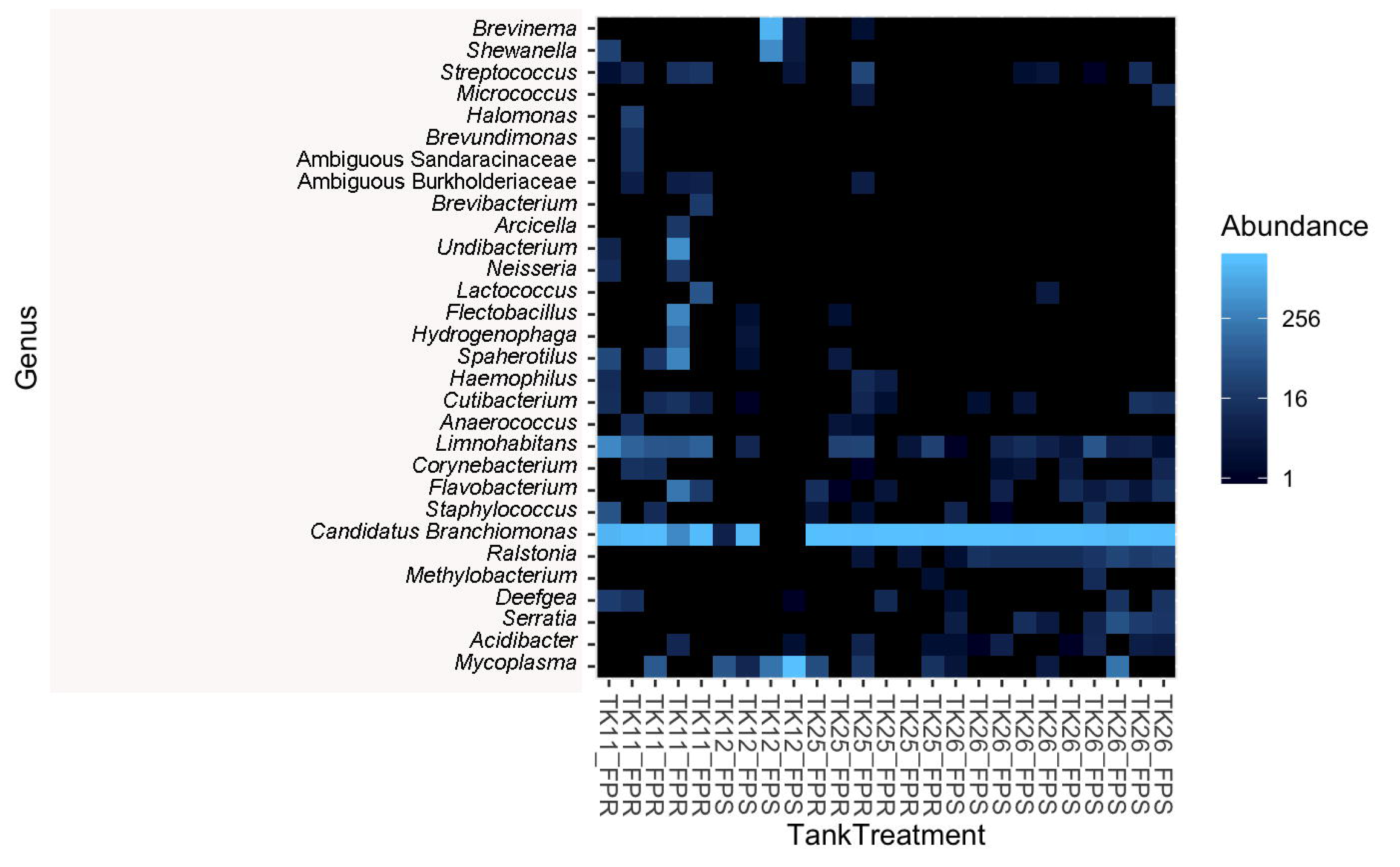
Heatmap representing the top 30 ASVs represented in the gill of ARS-Fp-R and ARS-Fp-S trout. Each column represents one individual fish. Each row represents one ASV.

### 3.7. Random Forest Modeling

Random forest modeling [31] was performed in order to see if a machine learning module could accurately predict treatment group as well as spleen index based on microbial community composition. In the mid-gut, this model was able to accurately predict 100% of FPR samples, but only 50% of FPS samples (Fig 7). Table 1 shows features at the genus level that were rendered as being important in classifying treatment group. Due to the lower sampling size in the gills, we did not include random forest analyses of the gill samples. Finally, we also performed a random forest regressor to attempt to predict spleen index based on gut microbial composition, and found that there was no correlation between these two variables.

**Fig 7.**
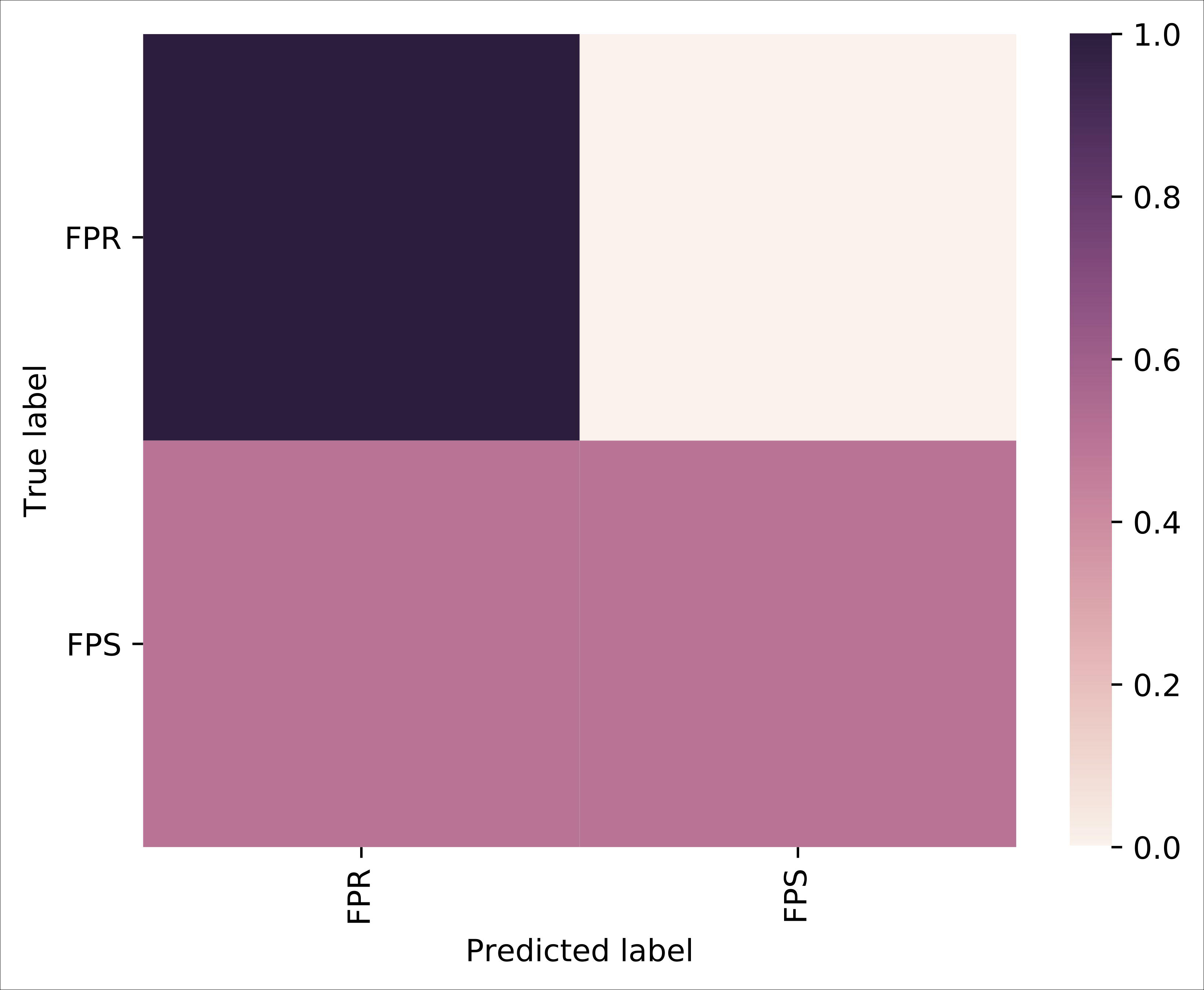
Confusion matrix for random forest modeling in the gut. This heatmap represents how frequently samples within each line were correctly classified. The correct label is represented on the y-axis and the predicted label is represented on the x-axis, with the prediction frequency shown as a gradient.

**Table 1:**
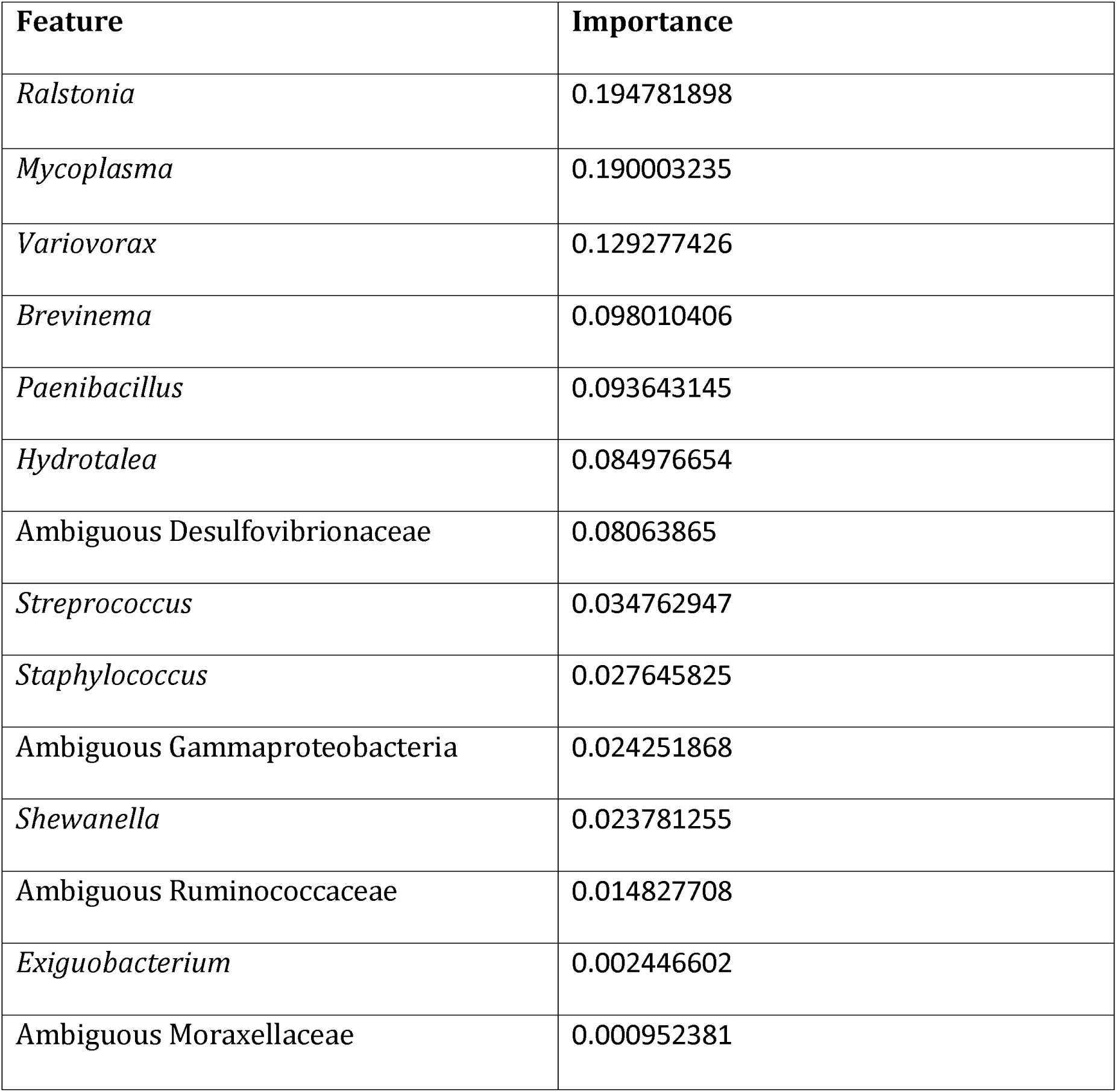
Features at the genus level that were identified by random forest modeling as being important in the classification of treatment group.

| Feature | Importance |
| --- | --- |
| *Ralstonia* | 0.194781898 |
| *Mycoplasma* | 0.190003235 |
| *Variovorax* | 0.129277426 |
| *Brevinema* | 0.098010406 |
| *Paenibacillus* | 0.093643145 |
| *Hydrotalea* | 0.084976654 |
| Ambiguous Desulfovibrionaceae | 0.08063865 |
| *Streptococcus* | 0.034762947 |
| *Staphylococcus* | 0.027645825 |
| Ambiguous Gammaproteobacteria | 0.024251868 |
| *Shewanella* | 0.023781255 |
| Ambiguous Ruminococcaceae | 0.014827708 |
| *Exiguobacterium* | 0.002446602 |
| Ambiguous Moraxellaceae | 0.000952381 |

## 4. Discussion

Commensal microbes have co-evolved with their eukaryotic counterparts, forming an intricate relationship that benefits both parties involved. Several studies have revealed that host genetics influences gut microbiota composition in a variety of species, including humans and rodents [4,5], chickens [32], and *Drosophila* [33]. However, other factors such as host diet and environmental factors are also deeply intertwined and deeply shape host microbial communities [7–9].

Teleost fish live in symbiosis with complex microbial communities that inhabit every mucosal barrier (gut, gills, skin and nose) [34]. Fish microbial community composition is influenced by age [35], tissue site [36], diet [37–39], stress [40] and pathogen infection [41]. However, few studies have investigated the impact host genetics have on shaping teleost microbiomes. A study on brook charr identified three quantitative trait loci associated with abundance of commensal strains in the skin [42]. Another study in Atlantic salmon found significant differences in the skin and gut microbial composition amongst distinct wild populations that are not likely attributed to environmental conditions alone [43]. These observations suggest that host genetics play an important role in teleost microbiome assembly, although more work is needed to better understand this relation.

Farmed fish are susceptible to many pathogens that threaten the sustainability of the finfish farming industry. Among the most prominent bacterial diseases, BCWD is particularly problematic in rainbow trout. The development of two genetic trout lines with different susceptibilities to BCWD agent, *F. psychrophilum*, offers an excellent platform to understand how host genetics may shape fish microbial communities. The current study suggests that host genetics influence the microbial community composition in the mid-gut, since the mid-gut bacterial community of the susceptible line was significantly more diverse than that of the resistant line. In agreement with previous studies, the mid-gut communities of both lines were dominated by *Mycoplasma sp*. This taxon appears to be highly abundant in the gastrointestinal microbiome of all salmonid species studied, including Atlantic salmon [44–46], rainbow trout [36,47] and Chinook salmon [48]. Interestingly, we identified a lower abundance of *Mycoplasma sp*. in the susceptible line, suggesting that disease susceptibility may be associated with decreased *Mycoplasma sp*. levels in the gut. In support, a recent study investigated the intestinal microbiome of offshore farmed Chinook salmon [48], finding that abundance of potential pathogenic *Vibrio sp*. appeared to be inversely correlated with the presence of *Mycoplasma sp*. *Mycoplasma sp*. are characterized by their uniquely small genome and lack of a cell wall, which makes culture-based approaches to studying this bacterium difficult to achieve. Considering the widespread presence and abundance of *Mycoplasma sp*. in salmonid gastrointestinal tract across a wide range of geographical locations, including both lab-reared fish and wild-caught, it appears as if there have been strong evolutionary forces that have enabled *Mycoplasma sp*. to thrive in this microenvironment. Future studies are needed to determine the nature of this relationship and the ability of *Mycoplasma sp*. to prevent pathogen colonization in the gastrointestinal tract of salmonids.

Potential opportunistic pathogens were observed within the mid-gut of the ARS-Fp-S line at a greater abundance than in the ARS-Fp-R line. Expansions of opportunistic taxa have been previously described in other fish microbiome studies including in the skin of Atlantic salmon experimentally infected with salmonid alphavirus (Reid, 2017) and in Atlantic salmon experimentally infected with the parasite *Lepeophtheirus salmonis* [41]. Specifically, we found *Brevinema sp*. and ambiguous members of the family Enterobacteriaceae in the susceptible line. *Brevinema sp*. was further identified as *Brevinema andersonii* at the species level in the Silva database, which is a pathogenic taxa that has been isolated in rodents [49]. However, upon sequence search using BLAST, they were most closely identified as an uncultured Spirochaeta sequence derived from an aquatic environment. Spirochaetes members have been described to cause disease in other marine species [50], and there are multiple well-studied members that are human pathogens [51]. Deeper taxonomic characterization would be necessary to determine if this ASV is in fact a pathogen in rainbow trout. Meanwhile, Enterobacteriaceae reads were most closely identified as *Hafnia paralvei*, which is associated with infections in a variety of animals, including fish [52]. Although the relative abundance of these taxa was small, these results suggest that ARS-Fp-R trout may possess a more resilient mid-gut microbiome, as compared to ARS-Fp-S. It is important to note that these fish were reared on spring water and sampled only at one timepoint. Additional studies are needed to examine whether these trends are observed in other environments and stages of development.

Differences in the microbial community composition between both trout lines were less pronounced in the gills compared to the gut, although it should be noted that the sample size was smaller in this tissue due to difficulties amplifying the 16S rDNA region in certain samples. Nonetheless, alpha diversity metrics were similar in each line. Interestingly, *Candidatus Branchiomonas* was the dominant taxon in both lines. This is a known pathogen that has been shown to cause gill epitheliocystis in Atlantic salmon [53], although it has not been previously described in rainbow trout. Fish from both lines were visually healthy, suggesting that *Candidatus Branchiomonas* may be a common member of the trout gill microbiome in certain environments. Further studies should evaluate which factors favor the colonization of *Candidatus Branchiomonas* in trout gills.

*Flavobacterium sp*. was identified in the gills of both lines, although sequence search using BLAST did not yield species level taxonomic resolution. Although this taxon was present at low abundance in both lines, it was surprising that we detected higher *Flavobacterium sp*. abundance in the gill microbiome of the resistant line compared to the susceptible line. Considering that the resistant and susceptible lines maintained the survival phenotype following the challenge experiment, our results indicate that susceptibility to *F. psychrophilum* infection is not due to increased abundance of this pathogen as a member of the indigenous microbial community. Disease susceptibility may be instead due to the greater ability of this pathogen to displace the microbial communities in the susceptible line compared to the resistant line.

One of the interesting aspects of the present study was the identification of differences in the microbial composition between tanks of the same trout genetic lines. All tanks were supplied water from the same source, and proper water quality was maintained throughout the experiment. This tank effect complicates the ability to discern between genetic lines, although there still appears to be notable differences when comparing each line, particularly in the gut as discussed earlier. There were similarities between two of the neighboring tanks that housed different lines, in tanks 11 and 12. However, this did not occur in the other two neighboring tanks, as the susceptible trout from tank 26 displayed an expansion of pathogenic taxa compared to tank 25, which housed resistant trout.

Work in zebrafish has shown that interhost dispersal can actually outweigh genetic factors that contribute to microbiome assembly [54]. In order to assess whether host genetics contribute to host microbiome assembly, all external factors that potentially contribute to this network must also be accounted for. In the present study, diet was consistent across fish of all tanks, and environmental factors such as water quality and temperature were also consistently maintained. Still, differences amongst tanks were present that add noise to the between-group comparisons. This finding highlights the importance of adequate experimental design in fish microbiome studies as well as the fact that different factors differentially shape microbial assembly at different tissue sites (i.e. gut versus gills).

## 5. Conclusions

In conclusion, the present study reveals differences in the microbial composition of the gut but not the gills of two rainbow trout lines with differential susceptibility to *F. psychrophilum* infection. Disease susceptibility was associated with a more diverse gut microbiome and the presence of potentially pathogenic taxa although important tank effects were also detected. Thus, selective breeding programs may not only select for host genetic factors but also, as a consequence, unique microbial assemblies, which in turn, may render the host more or less resilient to pathogen invasion or infection.

## Acknowledgements

We thank Kurt Schwalm and Dr. Darrell Dinwiddie for handling the Illumina MiSeq runs. We thank Dr. Timothy Leeds for breeding the ARS-Fp-R and ARS-Fp-S genetic lines and Travis Moreland for fish rearing and Jeremy Everson for assistance with fish sampling. This work was supported in part by Agricultural Research Service Project 1930-32000-006. Mention of trade names or commercial products in this publication is solely for the purpose of providing specific information and does not imply recommendation or endorsement by the U.S Department of Agriculture. USDA is an equal opportunity employer.

## Data Availability

All datasets obtained have been deposited at NCBI BioProject and are publicly available under BioProject ID number PRJNA488363.

